# Genetic manipulation of betta fish

**DOI:** 10.1101/2023.02.16.528733

**Authors:** Alec Palmiotti, Madison R Lichak, Pei-Yin Shih, Andres Bendesky

## Abstract

*Betta splendens*, also known as Siamese fighting fish or ‘betta’, are renowned for their astonishing morphological diversity and extreme aggressive behavior. Despite recent advances in our understanding of the genetics and neurobiology of betta, the lack of tools to manipulate their genome has hindered progress at functional and mechanistic levels. In this study, we outline the use of three genetic manipulation technologies, which we have optimized for use in betta: CRISPR/Cas9-mediated knockout, CRISPR/Cas9-mediated knockin, and Tol2-mediated transgenesis. We knocked out three genes: *alkal2l, bco1l*, and *mitfa*, and analyzed their effects on viability and pigmentation. Furthermore, we successfully knocked in a fluorescent protein into the *mitfa* locus, a proof-of-principle experiment of this powerful technology in betta. Finally, we used Tol2-mediated transgenesis to create fish with ubiquitous expression of GFP, and then developed a bicistronic plasmid with heart-specific expression of a red fluorescent protein to serve as a visible marker of successful transgenesis. Our work highlights the potential for the genetic manipulation of betta, providing valuable resources for the effective use of genetic tools in this animal model.

## 1 Introduction

The Siamese fighting fish (*Betta splendens*) or more commonly ‘betta’, is a species of freshwater fish known for its vibrant and diverse colors and fin morphologies, as well as its exceptional aggressive behavior. Native to Southeast Asia, betta have experienced a long history of domestication and selective breeding beginning at least 400 years ago (Kwon et al., 2022). Due to their morphological and phenotypic diversity, relatively compact genome size (∼440 Mb), their ease of growth in the laboratory, and their amenability to behavioral and neurobiological experimentation, betta have become an increasingly popular organism for scientific study (Lichak et al., 2022). The recent publication of high-quality reference genomes of both domesticated and wild betta has allowed for in-depth genetic and genomic analyses of sex determination, phenotypic traits such as pigmentation, fin shape and aggression, and of the evolutionary relationships between *Betta* species (Fan et al., 2018; Prost et al., 2020; Wang et al., 2021, 2022; Kwon et al., 2022; Yue et al., 2022; Zhang et al., 2022).

Establishing genetic tools in betta will advance their use as a powerful experimental system to study developmental processes and behavioral traits. Genetic manipulation of betta is facilitated by their reproductive biology: Betta fertilize externally and produce clutches of ∼250 eggs, each with a relatively large diameter of 1 mm (Valentin et al., 2015; Lichak et al., 2022). This enables the microinjection of zygotes in a similar manner to methods for genetic manipulation of zebrafish and medaka. Nevertheless, the asynchronous egg fertilization resulting from a protracted mating process that can last many hours, coupled with a short interval between fertilization and cell division, as well as a thick chorion, constitute significant challenges for genetic manipulation (Valentin et al., 2015; Lichak et al., 2022). Although a few studies have successfully utilized these tools, their reported success rates have been low, and none have reported germline transmission (Wang et al., 2021, 2022; Yue et al., 2022). Furthermore, although plasmid DNA microinjection has been used to make transgenic commercial betta (GloFish® Betta), the more controlled and efficient Tol2-based approach has not been performed. Tol2-mediated transgenesis has advantages over simple plasmid injections, such as improved rates of germline transmission and higher frequency of single copy integration (Kawakami, 2007).

In this study, we established effective protocols for three different genetic manipulation technologies in betta: CRISPR/Cas9 knockout, CRISPR/Cas9 knockin, and Tol2 transgenesis. We use CRISPR/Cas9 to knockout three genes, *ALK and LTK-ligand 2-like* (*alkal2l*), *beta-carotene oxygenase like-1* (*bco1l*), and *melanocyte inducing transcription factor a* (*mitfa*) and provide a phenotypic analysis of pigmentation of *alkal2l* germline crispants. We also report the first successful CRISPR/Cas9-mediated knockin, as well as the first use of Tol2-based transgenesis in betta. These methods, along with examples of their applications, highlight the potential for further adoption of genetic tools to study betta.

## 2 Results

### 2.1 CRISPR/Cas9 knockout editing of betta

We chose *alkal2l* and *bco1l* as targets for CRISPR/Cas9 editing based on a previous genetic mapping study from our lab that identified variation in these genes as likely contributing to blue or red coloration in ornamental betta (Kwon et al., 2022). *alkal2l* alleles present in blue fish are associated with an increase in the proportion of blue coloration covering the body, and a concomitant decrease in the proportion of red, whereas *bco1l* alleles likely modulate red hue (Kwon et al., 2022). Although the molecular mechanisms responsible for this phenotypic variation have not been studied in betta, *alkal2l* is necessary in zebrafish for the differentiation of iridophores that contribute to blue iridescence (Mo et al., 2017), and *bco1l* cleaves β-carotene, a red-orange molecule, into two molecules of all-trans retinal (Kwon et al., 2022). As a positive control of the induction of CRISPR/Cas9-mediated deletions, we also targeted a third gene, *mitfa*, using a guide RNA that has been validated in betta (Wang et al., 2021). We injected 1–4 celled embryos with a ribonucleoprotein (RNP) injection mix consisting of Cas9 protein and guide RNA (gRNA; **Figure 1A** and **Supplementary Figure S1**). We interrupted betta ∼2 hours after they began mating, as this maximizes the number of embryos at the 1–4 cell stage, and injected these embryos using a very sharp and beveled needle that facilitates the penetration of the thick chorion (**Supplementary Figure S2**). We then allowed surviving embryos to develop for 5 weeks before extracting their DNA from a fin clip and genotyping them using a T7 endonuclease I (T7EI) assay (**Supplementary Figure S3A**). Ten of the fourteen (71%) *alkal2l-*targeted fish for which PCR amplification of a genomic region surrounding the cleavage site was successful showed T7EI activity indicative of Cas9 cutting activity. The rate of CRISPR/Cas9-mediated mutagenesis was lower for *bco1l* (9/21; 43%), and lowest for *mitfa* (4/29; 14%), yet at sufficiently high rates to be a practical approach. (**Figure 1B**). Because betta *mitfa* knockout phenotypes have been reported previously (although only in the P_0_ generation), we focused on the *alkal2l* and *bco1l* mutants for further study.

**Figure 1.**
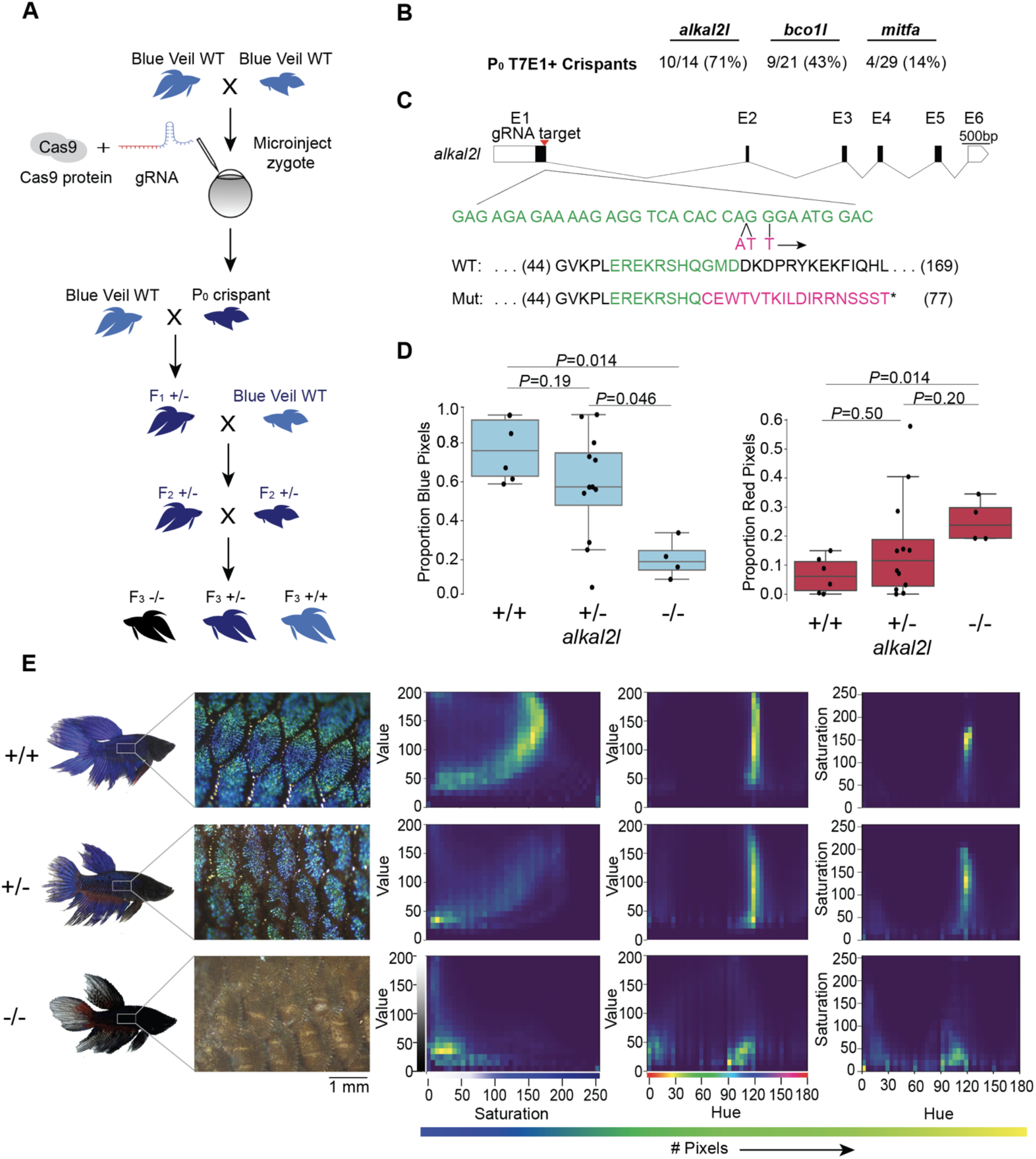
CRISPR/Cas9-mediated knockout in betta. **(A)** Experimental scheme for generating knockout betta. **(B)** Efficiency of knockout generation in P_0_, as determined by a T7EI assay. Numbers denote the fraction and percent of T7EI+ P_0_ individuals. **(C)** Schematic of *alkal2l* gene and the location of the gRNA target (top); wildtype DNA sequence in green and CRISPR/Cas9-induced mutations in magenta (middle); wildtype protein sequence (green shows the amino acids encoded by the wildtype sequence shown in the middle) and mutant sequence due to a frameshift (red shows the new mutant amino acids and * denotes a stop codon). **(D)** Boxplots showing proportion of blue (left) and red (right) pixels in *alkal2l* F_3_ crispants, according to genotype. *P-*values by Mann-Whitney test adjusted by Bonferroni correction. Boxes denote the interquartile range and whiskers the 5th and 95th percentiles, with a line at the median. **(E)** Representative images of betta individuals according to *alkal2l* genotype (left). Microscope images of the side of the body showing scales and the iridophore cover (center); the color of the microscope images appears different from the whole-body photographs on the left, due to the use of bright incident light under the microscope. Joint per-pixel quantification of hue, saturation and value by genotype (right).

We crossed the P_0_ crispants to wildtype betta of the same color and fin type as the parents of the P_0_ animals and genotyped the F_1_ offspring using a T7EI assay. We then sequenced the targeted loci of 15 T7EI+ F_1_ offspring (7 *bco1l*, 8 *alkal2l)* to identify the CRISPR-induced mutation. Sequence analysis revealed a variety of mutations in the F_1_ generations of *bco1l* and *alkal2l*. We chose one F_1_ crispant per gene that carried frameshift mutations leading to an early stop codon. We then crossed each of these F_1_ crispants in a second outcross to wildtype betta to obtain F_2_ generations containing heterozygotes genetically identical at the targeted loci (**Figure 1A**). For *bco1l*, this mutation was a T insertion that led to a frameshift coupled to a premature stop codon 21 amino acids downstream of the indel. For *alkal2l*, a two base-pair insertion plus a nearby single nucleotide mismatch led to a frameshift coupled to a premature stop codon 18 amino acids downstream of the indel (**Figure 1C**). We then crossed two F_2_ heterozygous sibling pairs to each other to obtain F_3_ animals, which we imaged to quantitatively analyze their coloration. Sequencing of the *bco1l* F_3_ individuals revealed that homozygous mutants were absent from the brood (19 +/+, 36 +/- and 0 -/-, *P*=0.0135 by Fisher’s exact test compared to the 1:2:1 Mendelian ratio), indicating that inheriting two mutant copies of *bco1l* is lethal. Post-imaging analyses of the color distributions and hues of *bco1l* +/+ vs +/- adult fish revealed no significant difference in the proportion of red and orange pixels, or of the joint hue, saturation and value (HSV) distributions (**Supplementary Figures S4A**,**B**). Therefore, allelic variation at *bco1l* between red and blue ornamental betta is unlikely to be composed of null alleles, and future reciprocal hemizygosity assays will evaluate the involvement of *bco1l* in the red-blue fish variation.

Next, we analyzed the impact of *alkal2l* in the F_3_ generation containing +/+, +/-, and -/- genotypes (**Supplementary Figure S3B**). We found a significant difference in the proportion of blue covering the body of *alkal2l* -/- compared to the other two genotypes, with -/- animals having only 25% as much blue cover as +/+ animals (**Figure 1D**). Heterozygotes were not different from wildtype, indicating that the *alkal2l* mutation is recessive (**Figure 1D**). The extent of blue iridescence in *alkal2l* -/- was reduced in both the body and the fins, yet the reduction was more prominent in the body, consistent with previous findings showing that *alkal2l* mRNA is present at 210× higher levels in the body compared to the fins (Kwon et al., 2022). Consistent with the reduction in blue coloration, the proportion of red surface was 5× more extensive in homozygous mutants compared to wild type fish (**Figure 1D**. Microscope images of the skin surface revealed that the blue iridescence mediated by iridophores is almost entirely absent on the body of *alkal2l* -/- animals, and is partially missing on the body of +/- mutants, suggesting that at least one functional *alkal2l* allele is crucial for iridescence in betta (**Figure 1E**). To better visualize differences in coloration, we plotted the joint distributions of per-pixel HSV (**Figure 1E**). The joint distributions of +/+ and +/- are similar, whereas -/- is an outlier. For hue vs saturation, in both +/+ and +/- animals, the majority of pixels have a hue value of ∼120 (corresponding to blue on a 180 degree hue color space), and these blue pixels range in saturation, with most falling between 100 and 200. The same graph for -/- shows a variety of different hues ranging from 0 to 120 with a saturation mostly below 50, indicating fewer blue pixels and less vibrant colors (**Figure 1E**). Furthermore, pixels in the -/- distributions are aggregated at much lower values than in the +/+ and +/- distributions, confirming that -/- mutants are darker and supporting the observation that knocking out *alkal2l* increases the visibility of deeper pigment layers that include melanophores. Together, these findings indicate that our method for obtaining CRISPR/Cas9 knockouts in betta is efficient and effective, that a homozygous knockout of *bco1l* is lethal, and that *alkal2l* is critical for blue coloration.

### 2.2 CRISPR/Cas9 Knock In

CRISPR/Cas9 knockin, which relies on the homology directed repair (HDR) pathway to integrate exogenous DNA into the host genome, has become the preferred method for the precise single-copy integration of various genome modifications such as fluorescent proteins, mutations relevant to specific disease models, and for epitope tagging (Hisano et al., 2015; Danner et al., 2017; Lino et al., 2018; Watakabe et al., 2018; Ranawakage et al., 2020). Recent advances in knockin technology, such as the use of 5’ biotinylated long homology arms, streptavidin-tagged Cas9, and *in vivo* linearization of the donor plasmid have been shown to increase the efficiency of knockins in zebrafish and mammalian cells (Gutierrez-Triana et al., 2018; Wierson et al., 2020). To develop knockin technology for betta, we adopted a recent efficient, cloning free CRISPR/Cas9 mediated knockin protocol that has been used in medaka (Seleit et al., 2021).

We targeted an insertion of GFP into *mitfa*, as this should lead to GFP expression in melanophores early in development, facilitating visual screening. To increase single-copy integration events and decrease the likelihood of donor DNA concatemerization (Gutierrez-Triana et al., 2018), we used primers with 5’ biotin modifications on the homology arms to amplify GFP by PCR (Seleit et al., 2021). We used monomeric streptavidin-tagged Cas9 mRNA (Cas9-mSA) to enhance Cas9 binding to the biotinylated donor constructs (Seleit et al., 2021). We then injected a mix consisting of *mitfa* gRNA (the same one we used for making knockouts), 5’-biotinylated donor DNA, and Cas9-mSA mRNA into 1-4 cell embryos (**Figure 2A**). As early as 24 hours post fertilization (hpf), robust GFP expression could be easily identified under the microscope in cells consistent with the location of neural crest cells, which give rise to melanophores (Silver et al., 2006), and at melanophores themselves. In three separate injections, the percentage of GFP+ fish were: 30%, 28%, and 26% (**Figure 2B**). At 2–3 days post fertilization (dpf), we sampled non-injected and GFP+ embryos for genotyping by PCR. We used primers for GFP, as well as ‘junction’ PCR primers, one of which binds *mitfa* and the other within GFP, to confirm GFP insertion at the intended locus. All 20 fish visibly expressing GFP were positive for GFP by PCR. 19/20 (95%) of these embryos were also positive for the junction PCR, indicating efficient integration at the *mitfa* locus in GFP+ embryos. Together, these findings indicate successful CRISPR/Cas9-mediated knockins in betta.

**Figure 2.**
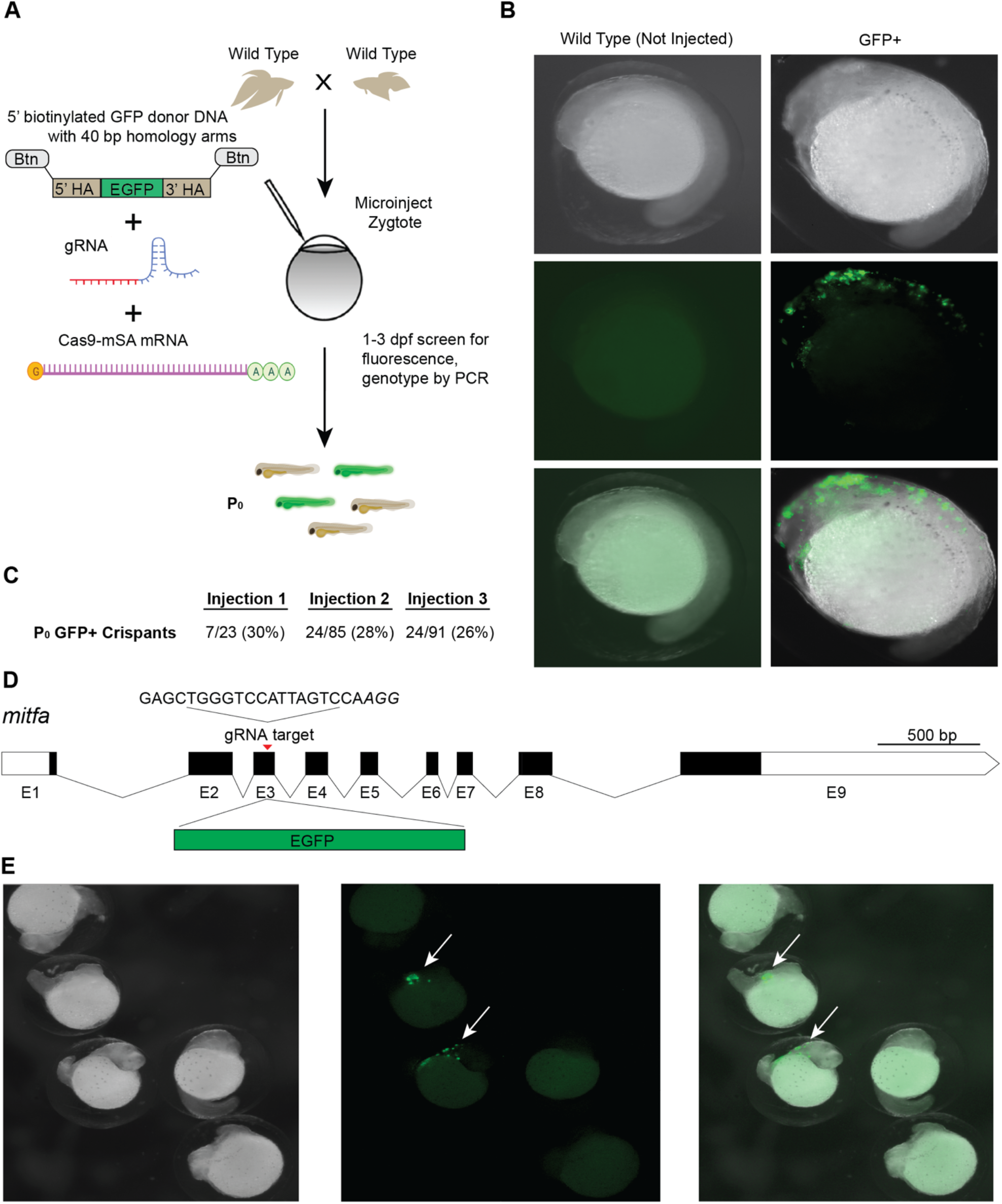
CRISPR/Cas9-mediated knockin in betta. **(A)** Experimental scheme for generating knockin betta. **(B)** Brightfield (top), fluorescence only (middle) and merged images of wild type (non-injected) betta and P_0_ GFP+ betta at 24 hpf. **(C)** Efficiency of knockin generation across injection of three different clutches. Numbers denote the fraction and percent of GFP+ individuals; 95% of these GFP+ animals are also positive for a GFP-targeted locus junction PCR (see Results). **(D)** Schematic of *mitfa* gene and location of the gRNA target for insertion of GFP. **(E)** Brightfield (left), fluorescence only (center) and merge (right) of 24-hpf injected embryos with clear GFP expression (arrow) and no expression.

### 2.3 Tol2-mediated transgenesis in betta

Tol2 transgenesis, which relies on the autonomous Tol2 transposable element first discovered in medaka (Kawakami et al., 1998), has become the predominant mechanism for transgenesis in zebrafish and in other vertebrates due to its high efficiency and simplicity of transgene construct design (Kawakami et al., 1998; Urasaki et al., 2006; Kawakami, 2007). The standard protocol for Tol2 transgenesis in vertebrates involves the co-injection of a plasmid that contains Tol2 transposon repeats flanking the DNA sequence to be randomly integrated in the genome, along with synthetic transposase mRNA. As a first step towards adopting Tol2 transgenesis in betta, we opted for a promoter that would drive ubiquitous expression of GFP, facilitating successful screening under the microscope. To that end, we cloned the 4.2 kb region upstream of the betta β-actin (*actb*) transcription start site in front of EGFP in a Tol2 plasmid to make pAB-1. We then injected a mix consisting of pAB-1 and transposase mRNA into 1–4 celled betta embryos (**Figure 3A**). In injections of six independent clutches, robust GFP expression indicative of transgenesis was apparent in the majority of embryos, with percentages ranging from 78% to 91% (**Figure 3E**). Expression was visible starting at 20 hpf. During the juvenile developmental period of 10–30 dpf, GFP expression was especially prominent in muscle, but was also visible in a variety of tissues including the brain, heart, bladder and intestine (**Figure 3B**). GFP expression remained visible into adulthood, suggestive of genomic integration rather than transient expression from the injected plasmid. To assess the integration of the transgene into the germline, we crossed five P_0_ transgenics with GFP expression to wild type betta. Of these five crosses, one generated offspring with GFP expression (26/153 larvae; 16%), indicating germline transmission of the transgenic construct (**Figure 3C**).

**Figure 3.**
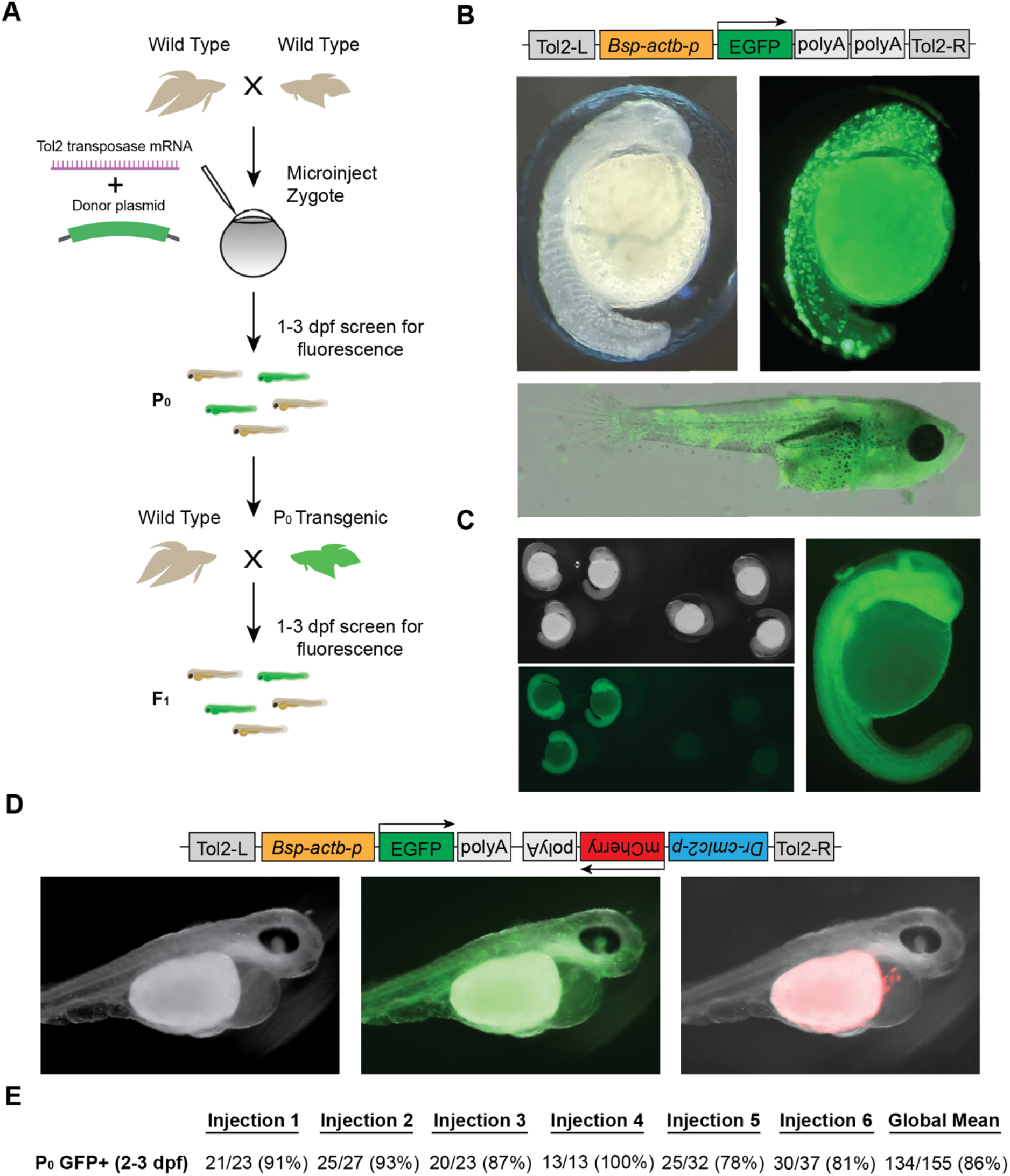
Tol2-mediated transgenesis in betta. **(A)** Experimental scheme for generating transgenic betta. **(B)** Schematic of pAB-1 *actb*:EGFP Tol2 plasmid (top), brightfield image of 24-hpf injected embryo (left) and fluorescent images of 24-hpf embryo (right) and ∼30-dpf betta (bottom). **(C)** Brightfield (top left) and fluorescent bottom left, right) images of *actb*:EGFP F_1_ transgenics. **(D)** Schematic of pAB-16 Tol2 bicistronic plasmid (top) and images of 3-dpf injected embryos using bright field (left) and GFP filter (middle) and mCherry filter (right). **(E)** Efficiency of transgenic generation across injections of six different clutches. Numbers denote the fraction and percentage of GFP+ P_0_ individuals.

Next, to make a bicistronic plasmid expressing both GFP ubiquitously, as well as mCherry in the heart to serve as an additional visible marker, we cloned the 268 bp region upstream of zebrafish *cmlc2* (Huang et al., 2003) in front of mCherry into pAB-1 to make pAB-16. We injected pAB-16 into 1-4 celled embryos using the same injection mix components as above. At 3 dpf, we observed both ubiquitous GFP expression, as well as robust mCherry expression in the heart of injected embryos (**Figure 3D**). At 3 dpf, but not at 1 and 2 dpf, mCherry expression in the heart was bright enough that it could be used to quickly screen for transgenesis. Together, these results indicate that Tol2-mediated transgenesis is a highly effective method for generating transgenics in betta. Our constructs also offer effective visual markers of transgenesis, and constitute useful starting points for cloning future transgenes.

## 3 Discussion

We established protocols for three genetic modification technologies in betta —CRISPR/Cas9 knockout, CRISPR/Cas9 knockin, and Tol2 transgenesis— each with efficiencies on par with current methodologies developed for more established model organisms such as zebrafish and medaka. CRISPR/Cas9 knockout has been previously used to study betta, yet our P_0_ mutagenesis efficiencies for the three genes we targeted (14%, 43%, and 71%) are markedly higher than the efficiency reported by others (4/2000; 0.2%) when targeting *dmrt1* in betta (Wang et al., 2022). CRISPR mutagenesis efficiencies using our methods are comparable to those seen in zebrafish, where single gRNA editing efficiencies initially ranged from ∼25-60% (Hwang et al., 2013; Liu et al., 2019), and more recent reports using multi-guide RNP complexes report editing efficiencies ranging from 80-100% (Klatt Shaw and Mokalled, 2021; Kroll et al., 2021). Furthermore, our P_0_ knockin targeting efficiency (28%) is consistent with efficiencies in medaka (11-59%) using the same method (Seleit et al., 2021), and our targeting efficiency for making transgenics using the Tol2 transposon system (86%) is close to that of zebrafish (∼100%) (Urasaki et al., 2006). Thus, the methods we provide constitute powerful tools to manipulate the betta genome. These tools will empower modern approaches to study the genetic mechanisms underlying the astonishing morphological diversity of betta, as well as its unique aggressive behavior.

## 4 Materials and Methods

### 4.1 Husbandry

Animals were maintained according to our standard procedures (Lichak et al., 2022). All animal protocols were approved by the Columbia University Institutional Animal Care and Use Committee. The night prior to injections, we set up mating pairs according to published protocols (Lichak et al., 2022). We added each mating pair to a 5 L acrylic tank filled to a height of 10 cm with system water. Each tank contained one terra cotta pot, one acrylic plant, and a floating piece of dried Indian almond leaf (*Terminalia catappa*, from SunGrow) to facilitate bubble nest building and allow for convenient egg collection. During mating, tanks were kept at 28–30 °C using heating mats (VivoSun, Ontario, CA), and were covered with acrylic to maintain high levels of humidity (∼90%) relative to the betta facility (∼25%). The following morning, we checked if fish were mating, and allowed for 1–3 hours of undisturbed mating before collecting eggs for microinjection. Eggs were then collected by carefully inverting the almond leaf containing the bubble nest and then gently washing the nest containing the eggs into a 250 mL beaker filled with system water.

### 4.2 Microinjection

Prior to the day of injection, we made microinjection needles and injection molds, and prepared working concentrations of components of microinjection mixes. To make microinjection needles we used a micropipette puller (Model P-1000, Sutter Instrument, Novato, CA) to pull 4-inch long thin wall filamented glass capillary tubes (TW100F-4, World Precision Instruments, Sarasota, FL) into injection needles using custom settings: Heat: 480, Pull: 90, Velocity: 80, Delay: 120, Pressure: 230, Ramp: 467. We then used micro-scissors (15000-08, Fine Science Tools, Foster City, CA) to cut the pulled needles at a ∼45° angle under a dissection microscope so as to bevel the needle to facilitate piercing of the thick chorion (**Supplementary Figure S2**). We then stored the pulled needles in a 150 mm petri dish on mounting putty (XTREME Putty, Tombow, Suwanee, GA) until use. We made injection molds to hold embryos for injection by dissolving 1 g agarose in 50 mL 1× E3 medium containing 0.0005% methylene blue, pouring the hot agarose solution into a petri dish, then placing the dry injection mold (TU-1, Adaptive Science Tools, Worcester, MA) into the solution when it reached 55°C. A 60× solution of E3 medium can be made by combining 8.7 g NaCl, 0.4 g KCl, 1.45 g CaCl_2_·2H_2_O, 2.45 g MgCl_2_·6H_2_O and bringing to 500 mL using reverse osmosis water, followed by pH adjustment to 7.2 using NaOH. We removed the mold when the agarose solution solidified and stored the plate containing the mold at 4°C.

In advance of injections, we also resuspended CRISPR RNAs (crRNAs) (Alt-R® CRISPR-Cas9 crRNA XT, 2 nmol, Integrated DNA Technologies, Coralville, IA) and tracrRNA (1072532, Alt-R® CRISPR-Cas9 tracrRNA, 5 nmol, Integrated DNA Technologies) to a working concentration of 100 µM using TE buffer (1 mM EDTA, 10 mM Tris-Cl; pH 8.0). We then assembled the gRNA on ice by combining 1.5 µL 100 µM crRNA, 1.5 µL 100 µM tracrRNA, and 47 µl Duplex buffer (11-05-01-03, Integrated DNA Technologies), and then heating the solution to 95 °C for 5 minutes before removing from heat and letting the solution cool passively to room temperature (21-22° C) on benchtop. These were stored at −70 °C until injection. We also diluted Cas9 protein (1081058, Alt-R® S.p. Cas9 Nuclease V3, 10 µg/µL, Integrated DNA Technologies) by combining 0.5 µL stock Cas9 protein with 9.5 µL Cas9 buffer (20 mM HEPES, 150 mM KCl, pH 7.5) to a final concentration of 500 ng/µL. This was stored at −20 °C until injection. On the day of injection, while fish mated and prior to collecting eggs, we prepared the RNP microinjection mix. For CRISPR/Cas9 knockouts we combined 2 µL 3 µM gRNA and 2 µL 500 ng/µL Cas9 protein and incubated at 37°C for 10 minutes before placing on ice and adding ∼0.2 µL 0.05% phenol red for visualization while injecting.

The CRISPR/Cas9 knockin microinjection mix consisted of 2 µl 3 µM gRNA, 1.5 µL 100 ng/µL 5’ biotinylated donor DNA, 1.5 µL 400 ng/µL Cas9-mSA mRNA, and ∼0.2 µL 0.05% phenol red. The gRNA was assembled in the same manner as described above for our knockouts. The 5’ biotinylated donor DNA was made by amplifying GFP from a plasmid using primers with biotinylated 40 bp overhangs that annealed to the host genome surrounding the cleavage site in frame. We confirmed the expected size of the PCR product by running an aliquot on a gel, and then used the DNA Clean and Concentrator Kit (D4209, Zymo Research, Irvine, CA) to purify the PCR reaction before diluting to a final concentration of 100 ng/µL in nuclease free water. To prepare Cas9-mSA-mRNA, we linearized PCS2+Cas9-mSA using NotI-HF (R3189S, New England Biolabs, Waltham, MA) and then transcribed it using mMessage mMachine SP6 (AM1340, Invitrogen, Waltham, MA). We then purified mRNA using the RNA clean and concentrator kit (R1017, Zymo Research), quantified it using a NanoDrop One (13-400-519, Thermo Fisher Scientific, Waltham MA), and made 400 ng/µL aliquots, which we stored at −70 °C until use.

The Tol2 transgenesis microinjection mix consisted of 0.5 µL 250 ng/µL plasmid DNA, 0.5 µL 250 ng/µL transposase mRNA, 2.5µL 0.4M KCl, 1 µL ddH_2_O, ∼0.2 µL 0.05% phenol red. To prepare Tol2 transposase mRNA, we linearized pCS2-zT2TP (a gift from Jamie Gagnon) using NotI-HF and then transcribed it using mMessage mMachine SP6. We then purified mRNA using the RNA clean and concentrator kit and made 250 ng/µL aliquots quantified using the NanoDrop One, which we stored at −70 °C until use.

After collecting the eggs from the mating tanks, we used a glass pipette to transfer eggs from the 250 mL beaker to our microinjection mold. To prevent the eggs from moving around in the mold, we removed most of the water leaving just enough to prevent the eggs from drying. We then loaded 1 µl of the microinjection mix into the beveled needle using a P2.5 pipette by slowly adding the mix to the back of the needle and then allowing capillary action to bring the mix to the needle tip. After ensuring the micromanipulator (M-152, Narishige, Setagaya, Tokyo) was in the proper position (able to move freely in all directions), we loaded the needle into the mount of the micromanipulator. After confirming most embryos were not past the 1-4 cell stage, we microinjected ∼1 nL microinjection mix into each embryo (either the single cell or one of the dividing cells) using a FemtoJet 4i microinjector (Eppendorf, Hamburg, Germany) (**Supplementary Figure S1**). After each round of injections, we used a plastic squeeze bottle filled with 1× E3 medium to gently wash the injected embryos into a glass petri dish, and then placed the petri dish in a 28 °C incubator.

### 4.3 Imaging

We photographed sexually mature fish inside a plastic tank using a Canon EOS RP with a Macro 100 mm lens (Macro Lens EF 100mm, Canon, Tokyo, Japan), as described (Kwon et al., 2022). All photos were taken inside a photo studio tent with white light emitting diodes. In each photo, a color card and ruler were visible, and a mirror was placed on the outside of the tank facing the fish to facilitate flaring, allowing for visualization of fins and consistency amongst photos. Before each imaging session, we calibrated the camera using the white balance (CT24-23-1424) on the 24ColorCard (CameraTrax).

### 4.4 Color Analysis

We used a custom Python script (**Supplementary Code**) to automatically threshold each image and determine the hue, saturation, and value (HSV) of each pixel of the body. We then calculated the joint distributions of HSV and the fraction of blue and red pixels. On a 0–180 hue scale, we defined blue as the range from 110–130 and red as the range from 0–30 (see hue scale bar in **Figure 1E**). For the *bco1l* color analysis, we differentiated orange and red pixels by defining red as the range from 0– 15, and orange as the range from 15–30.

### 4.5 Genotyping

We used various methods for genotyping crispants. P_0_, F_1_, and F_2_ crispants were fin clipped at 5 weeks and genotyped by a T7EI assay (M0302, New England Biolabs). We extracted DNA using the Quick-DNA Microprep Kit (D3021, Zymo Research). We Sanger sequenced *alkal2l* and *bco1l* F_1s_ and used CRISP-ID (Dehairs et al., 2016) to aid with identifying indels. To genotype *alkal2l* +/- × +/- crosses of the allele consisting of a 2-bp insertion plus a 1-bp mismatch, we used discriminative primers (**Supplementary Figure S3B**) annealing specifically to mutant and wild type alleles. To increase specificity of these primers to their respective alleles, we introduced an extra mismatch (Chen and Schedl, 2021). To genotype the *bco1l* +/- × +/- crosses of the 1-bp insertion allele, we Sanger sequenced the progeny. To genotype P_0_ knockin crispants, we extracted DNA from 2–4 dpf whole larvae using the Quick-DNA Microprep Kit (D3021, Zymo Research). We then performed a PCR for GFP as well as a ‘junction PCR’ to amplify a region including both *mitfa* and GFP to ensure integration at the intended locus.

### 4.6 Cloning

To clone the Tol2-based transgenesis plasmids, we first amplified the 4,165 bp region upstream of the transcription start site of the betta *actb* gene using Q5 polymerase (M0491S, New England Biolabs) from an ornamental betta. We then replaced the ubiquitin promoter in the pDestTol2pA2_ubi:EGFP plasmid (27323, Addgene; Mosimann et al., 2011) with the amplified *actb* promoter region using bacterial *in vivo* assembly (García-Nafría et al., 2016). We then purified plasmids for injection using the ZymoPURE plasmid miniprep kit (D4210, Zymo Research). We fully sequenced all plasmids prior to injection to confirm intended sequences. To create the bicistronic plasmid, we first linearized *actb*:EGFP and then amplified a 1.3 kb region from pBH mcs (a gift from Michael Nonet) containing the 268 bp of the *cmlc2* promoter region, mCherry, and the polyadenylation and large T antigen signals. We then carried out all subsequent steps as described above.

### 4.7 Primer Sequences

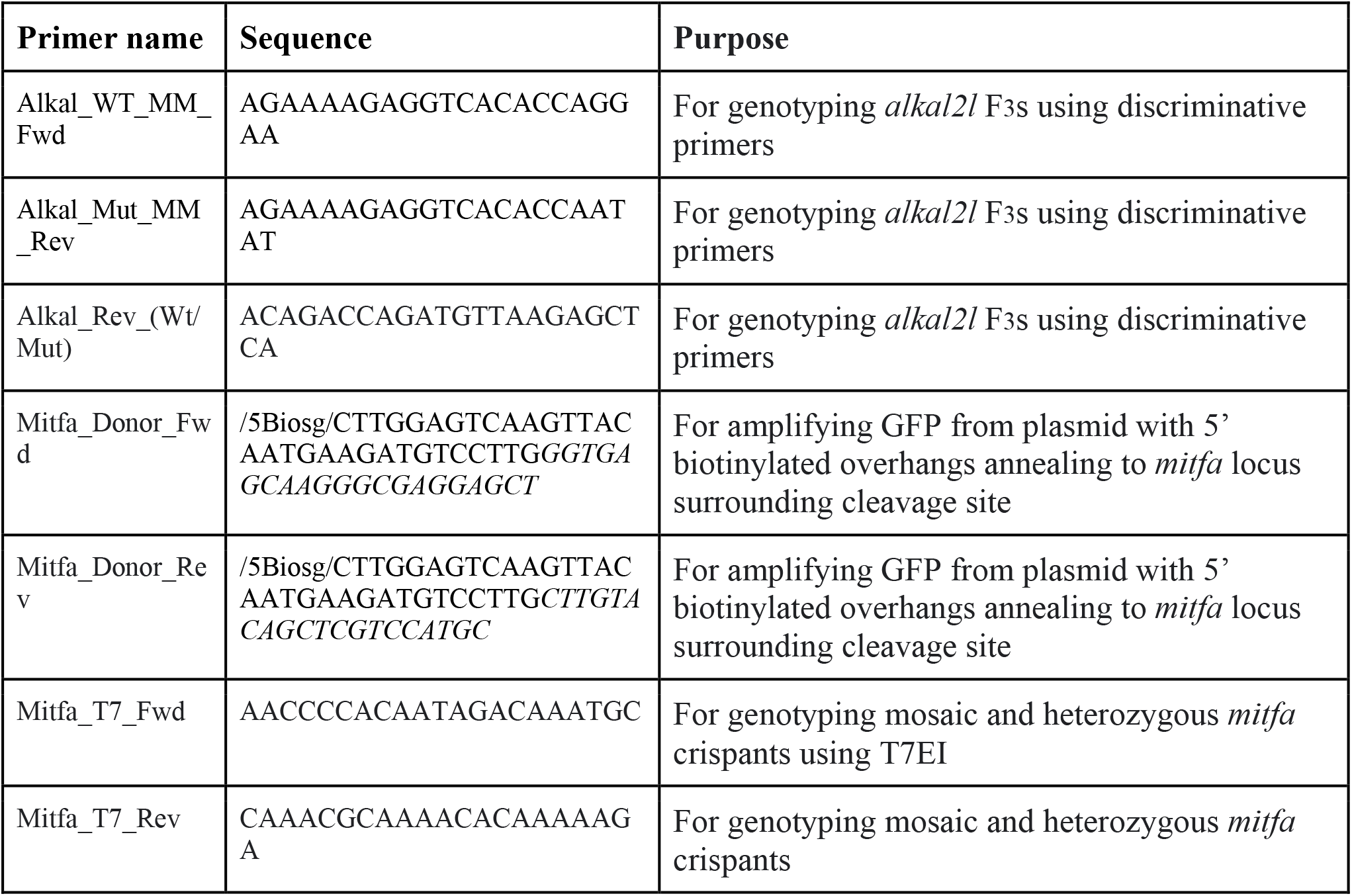
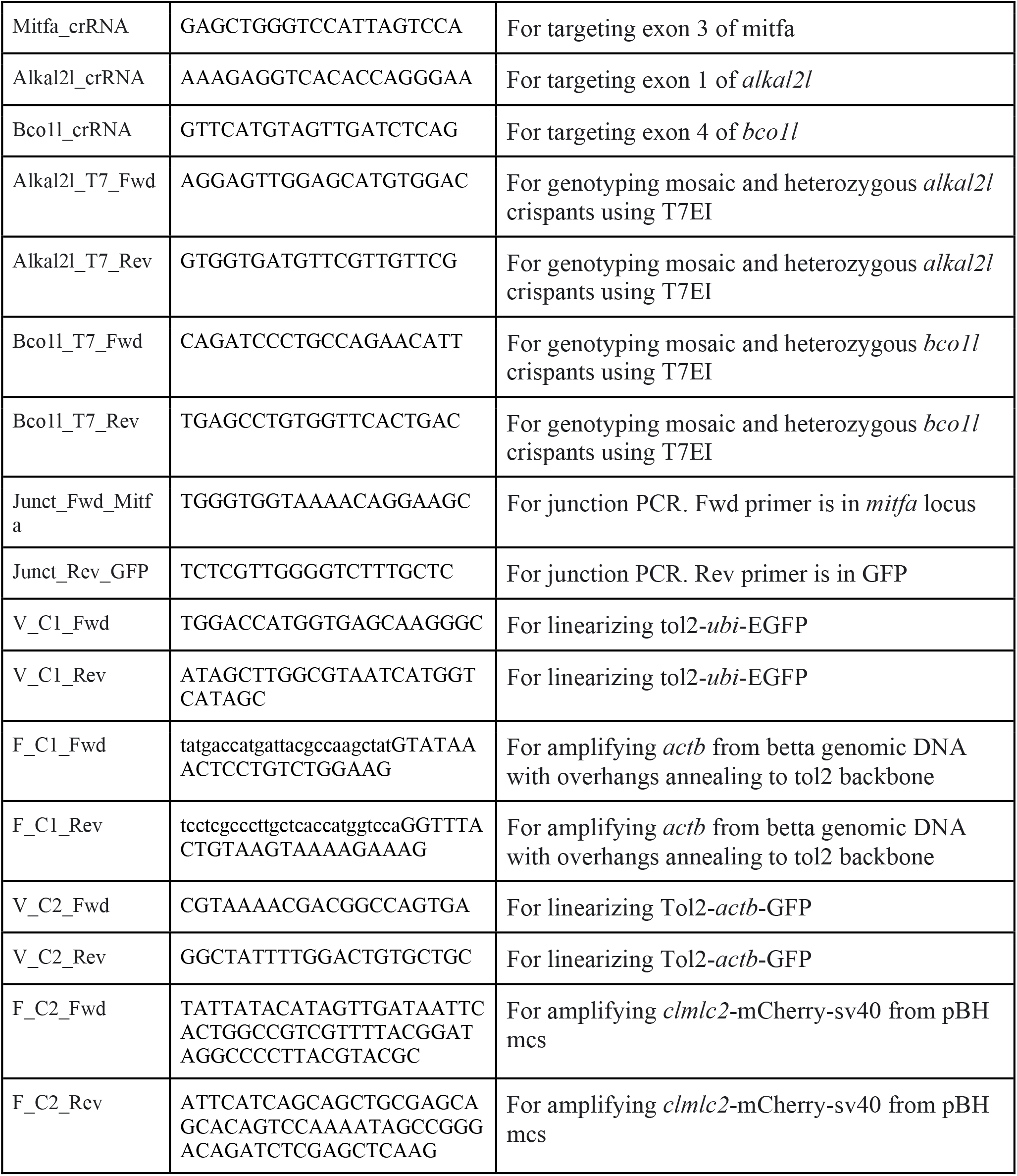

## 5 Author Contributions

AP and MRL contributed to conceptualization of the research project, investigation, data curation, methodology development and data analysis. AB and P-YS contributed to conceptualization of the research project, methodology development and supervision. AP (original draft) and AB wrote the manuscript. All authors contributed to the review and editing of the manuscript and approved the submitted version.

## 8 Acknowledgements

Michael Nonet and Jamie Gagnon shared reagents. David Ng provided technical assistance. Joshua Barber provided advice for microinjections and fish care.

## Bibliography

Chen, J., and Schedl, T. (2021). A simple one-step PCR assay for SNP detection. MicroPublication Biol. 2021. doi: 10.17912/micropub.biology.000399.

Danner, E., Bashir, S., Yumlu, S., Wurst, W., Wefers, B., and Kühn, R. (2017). Control of gene editing by manipulation of DNA repair mechanisms. Mamm. Genome Off. J. Int. Mamm. Genome Soc. 28, 262–274. doi: 10.1007/s00335-017-9688-5.

Dehairs, J., Talebi, A., Cherifi, Y., and Swinnen, J. V. (2016). CRISP-ID: decoding CRISPR mediated indels by Sanger sequencing. Sci. Rep. 6, 28973. doi: 10.1038/srep28973.

Fan, G., Chan, J., Ma, K., Yang, B., Zhang, H., Yang, X., et al. (2018). Chromosome-level reference genome of the Siamese fighting fish Betta splendens, a model species for the study of aggression. GigaScience 7, giy087. doi: 10.1093/gigascience/giy087.

García-Nafría, J., Watson, J. F., and Greger, I. H. (2016). IVA cloning: A single-tube universal cloning system exploiting bacterial In Vivo Assembly. Sci. Rep. 6, 27459. doi: 10.1038/srep27459.

Gutierrez-Triana, J. A., Tavhelidse, T., Thumberger, T., Thomas, I., Wittbrodt, B., Kellner, T., et al. (2018). Efficient single-copy HDR by 5’ modified long dsDNA donors. eLife 7, e39468. doi: 10.7554/eLife.39468.

Hisano, Y., Sakuma, T., Nakade, S., Ohga, R., Ota, S., Okamoto, H., et al. (2015). Precise in-frame integration of exogenous DNA mediated by CRISPR/Cas9 system in zebrafish. Sci. Rep. 5, 8841. doi: 10.1038/srep08841.

Hwang, W. Y., Fu, Y., Reyon, D., Maeder, M. L., Tsai, S. Q., Sander, J. D., et al. (2013). Efficient genome editing in zebrafish using a CRISPR-Cas system. Nat. Biotechnol. 31, 227–229. doi: 10.1038/nbt.2501.

Kawakami, K. (2007). Tol2: a versatile gene transfer vector in vertebrates. Genome Biol. 8 Suppl 1, S7. doi: 10.1186/gb-2007-8-s1-s7.

Kawakami, K., Koga, A., Hori, H., and Shima, A. (1998). Excision of the tol2 transposable element of the medaka fish, Oryzias latipes, in zebrafish, Danio rerio. Gene 225, 17–22. doi: 10.1016/s0378-1119(98)00537-x.

Klatt Shaw, D., and Mokalled, M. H. (2021). Efficient CRISPR/Cas9 mutagenesis for neurobehavioral screening in adult zebrafish. G3 11, jkab089. doi: 10.1093/g3journal/jkab089.

Kroll, F., Powell, G. T., Ghosh, M., Gestri, G., Antinucci, P., Hearn, T. J., et al. (2021). A simple and effective F0 knockout method for rapid screening of behaviour and other complex phenotypes. eLife 10, e59683. doi: 10.7554/eLife.59683.

Kwon, Y. M., Vranken, N., Hoge, C., Lichak, M. R., Norovich, A. L., Francis, K. X., et al. (2022). Genomic consequences of domestication of the Siamese fighting fish. Sci. Adv. 8, eabm4950. doi: 10.1126/sciadv.abm4950.

Lichak, M. R., Barber, J. R., Kwon, Y. M., Francis, K. X., and Bendesky, A. (2022). Care and use of Siamese fighting fish (Betta splendens) for research. Comp. Med. 72, 169–180. doi: 10.30802/AALAS-CM-22-000051.

Lino, C. A., Harper, J. C., Carney, J. P., and Timlin, J. A. (2018). Delivering CRISPR: a review of the challenges and approaches. Drug Deliv. 25, 1234–1257. doi: 10.1080/10717544.2018.1474964.

Liu, K., Petree, C., Requena, T., Varshney, P., and Varshney, G. K. (2019). Expanding the CRISPR toolbox in zebrafish for studying development and disease. Front. Cell Dev. Biol. 7, 13. doi: 10.3389/fcell.2019.00013.

Mo, E. S., Cheng, Q., Reshetnyak, A. V., Schlessinger, J., and Nicoli, S. (2017). Alk and Ltk ligands are essential for iridophore development in zebrafish mediated by the receptor tyrosine kinase Ltk. Proc. Natl. Acad. Sci. U. S. A. 114, 12027–12032. doi: 10.1073/pnas.1710254114.

Mosimann, C., Kaufman, C. K., Li, P., Pugach, E. K., Tamplin, O. J., and Zon, L. I. (2011). Ubiquitous transgene expression and Cre-based recombination driven by the ubiquitin promoter in zebrafish. Dev. Camb. Engl. 138, 169–177. doi: 10.1242/dev.059345.

Prost, S., Petersen, M., Grethlein, M., Hahn, S. J., Kuschik-Maczollek, N., Olesiuk, M. E., et al. (2020). Improving the chromosome-level genome assembly of the Siamese fighting fish (Betta splendens) in a university master’s course. G3 10, 2179–2183. doi: 10.1534/g3.120.401205.

Ranawakage, D. C., Okada, K., Sugio, K., Kawaguchi, Y., Kuninobu-Bonkohara, Y., Takada, T., et al. (2020). Efficient CRISPR-Cas9-mediated knock-in of composite tags in zebrafish using long ssDNA as a donor. Front. Cell Dev. Biol. 8, 598634. doi: 10.3389/fcell.2020.598634.

Seleit, A., Aulehla, A., and Paix, A. (2021). Endogenous protein tagging in medaka using a simplified CRISPR/Cas9 knock-in approach. eLife 10, e75050. doi: 10.7554/eLife.75050.

Silver, D. L., Hou, L., and Pavan, W. J. (2006). The genetic regulation of pigment cell development. Adv. Exp. Med. Biol. 589, 155–169. doi: 10.1007/978-0-387-46954-6_9.

Urasaki, A., Morvan, G., and Kawakami, K. (2006). Functional dissection of the Tol2 transposable element identified the minimal cis-sequence and a highly repetitive sequence in the subterminal region essential for transposition. Genetics 174, 639–649. doi: 10.1534/genetics.106.060244.

Valentin, F. N., do Nascimento, N. F., da Silva, R. C., Fernandes, J. B. K., Giannecchini, L. G., and Nakaghi, L. S. O. (2015). Early development of Betta splendens under stereomicroscopy and scanning electron microscopy. Zygote Camb. Engl. 23, 247–256. doi: 10.1017/S0967199413000488.

Wang, L., Sun, F., Wan, Z. Y., Yang, Z., Tay, Y. X., Lee, M., et al. (2022). Transposon-induced epigenetic silencing in the X chromosome as a novel form of dmrt1 expression regulation during sex determination in the fighting fish. BMC Biol. 20, 5. doi: 10.1186/s12915-021-01205-y.

Wang, L., Sun, F., Wan, Z. Y., Ye, B., Wen, Y., Liu, H., et al. (2021). Genomic basis of striking fin shapes and colors in the fighting fish. Mol. Biol. Evol. 38, 3383–3396. doi: 10.1093/molbev/msab110.

Watakabe, I., Hashimoto, H., Kimura, Y., Yokoi, S., Naruse, K., and Higashijima, S.-I. (2018). Highly efficient generation of knock-in transgenic medaka by CRISPR/Cas9-mediated genome engineering. Zool. Lett. 4, 3. doi: 10.1186/s40851-017-0086-3.

Wierson, W. A., Welker, J. M., Almeida, M. P., Mann, C. M., Webster, D. A., Torrie, M. E., et al. (2020). Efficient targeted integration directed by short homology in zebrafish and mammalian cells. eLife 9, e53968. doi: 10.7554/eLife.53968.

Yue, G. H., Wang, L., Sun, F., Yang, Z., Shen, Y., Meng, Z., et al. (2022). The ornamental fighting fish is the next model organism for genetic studies. Rev. Aquac. 14, 1966–1977. doi: 10.1111/raq.12681.

Zhang, W., Wang, H., Brandt, D. Y. C., Hu, B., Sheng, J., Wang, M., et al. (2022). The genetic architecture of phenotypic diversity in the Betta fish (Betta splendens). Sci. Adv. 8, eabm4955. doi: 10.1126/sciadv.abm4955.

