## Supplement for "Genetic manipulation of betta fish"

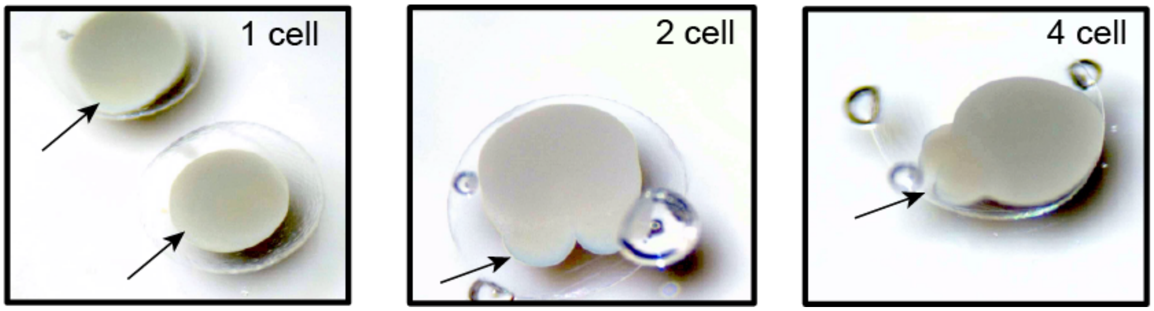


**Supplementary Figure 1.** Images of single-celled embryo (left), two-celled embryo (middle) and four-celled embryo (right)

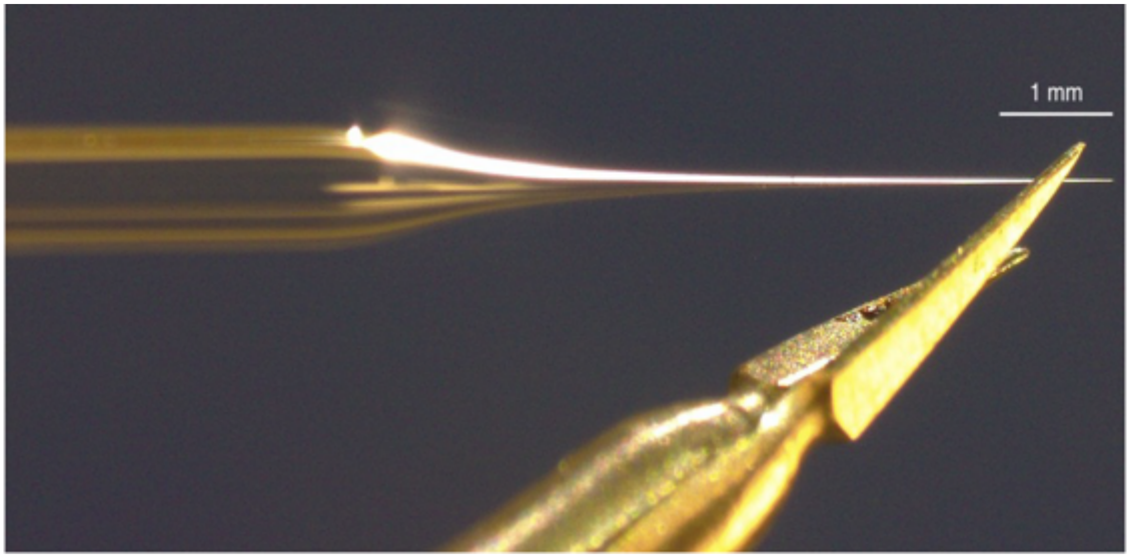


**Supplementary Figure 2.** Photograph of injection needle being cut ~0.5 mm from the tip at ~45º

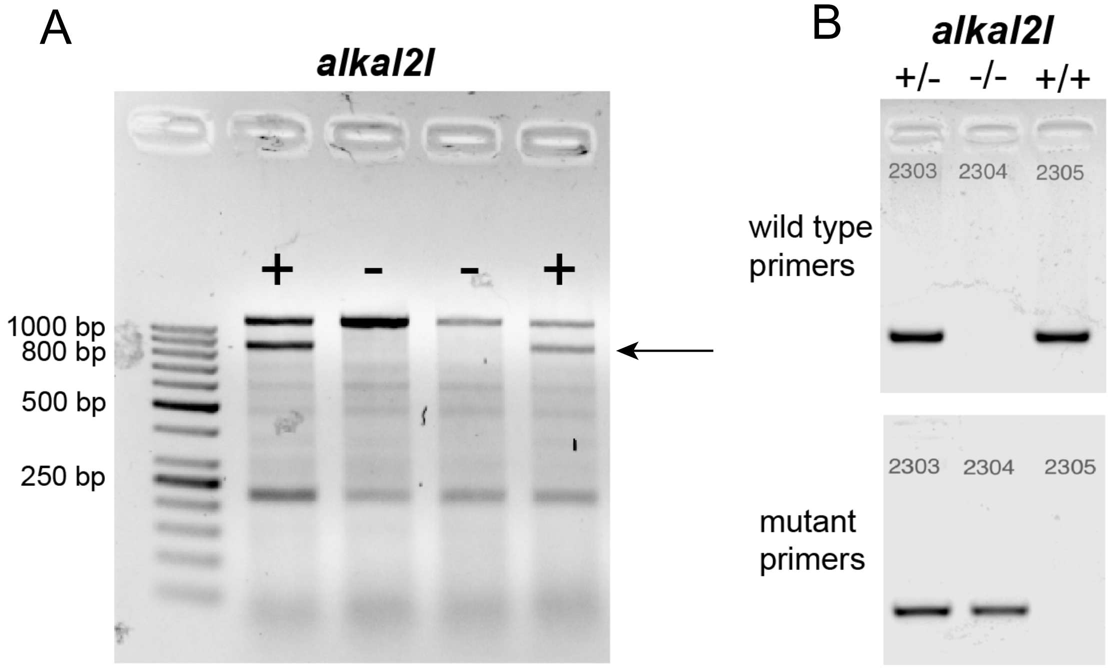


**Supplementary Figure 3. (A)** Representative T7EI assay for genotyping mosaic and heterozygous *alkal2l* crispants. Arrow points to extra band indicative of positive T7EI assay. **(B)** Representative discriminative primer assay for genotyping $F_3$ *alkal2l* +/+, +/- and -/- animals. 2303, 2304 and 2305 are fish unique IDs.

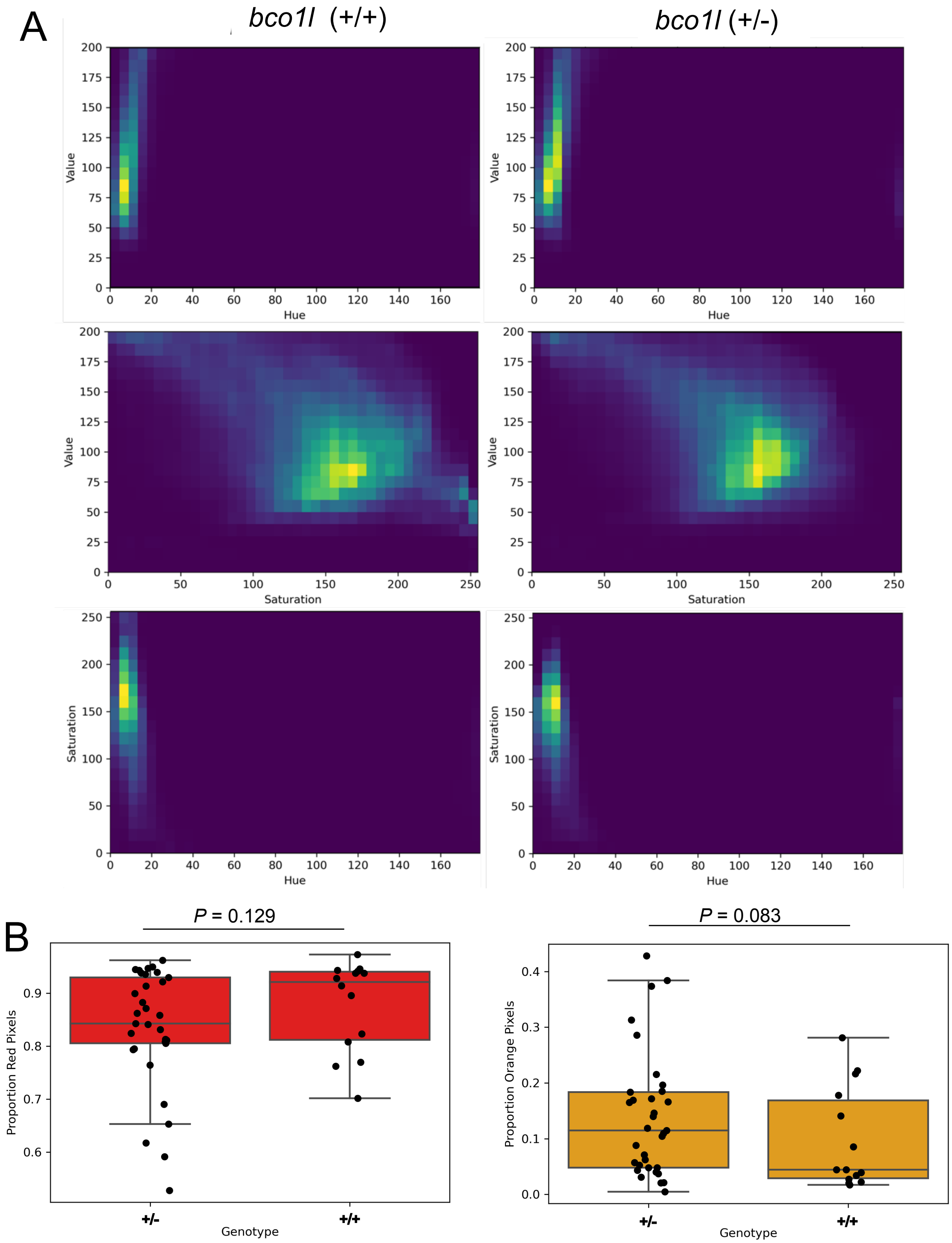


**Supplementary Figure 4. (A)** Joint HSV distributions of *bco1l* (+/+) and (+/-) animals **(B)** Boxplots showing proportion of orange (left) and red (right) pixels in *bco1l* crispants, according to genotype. *P* values by Mann-Whitney test adjusted by Bonferroni correction. Boxes denote the interquartile range and whiskers the 5th and 95th percentiles, with a line at the median.

**Supplementary Code.** *alkal2l* color analysis python3 code

```
import os
from glob import glob
import cv2 as cv
import numpy as np
import pandas as pd
from matplotlib import pyplot as plt
import seaborn as sns

img_fh = '/directory/2427_Homo_C1.tiff'
img = cv.imread(img_fh,cv.IMREAD_COLOR)
img_hsv = cv.cvtColor(img, cv.COLOR_BGR2HSV)
plt.imshow(cv.cvtColor(img_hsv, cv.COLOR_HSV2RGB))
plt.show()
hue_hist = cv.calcHist([img_hsv],[0],None,[256],[0,256])
sat_hist = cv.calcHist([img_hsv],[1],None,[256],[0,256])
val_hist = cv.calcHist([img_hsv],[2],None,[256],[0,256])
plt.plot(hue_hist)
plt.plot(sat_hist)
plt.plot(val_hist)
plt.legend(['hue','sat','value'])
plt.show()
mask = cv.inRange(img_hsv, np.array([0,0,0]), np.array([255,255,200]))
plt.imshow(mask)
plt.show()
res = cv.bitwise_and(img_hsv, img_hsv, mask=mask)
plt.imshow(cv.cvtColor(res, cv.COLOR_HSV2RGB))
plt.show()
hue_hist = cv.calcHist([img_hsv],[0],mask,[256],[0,256])
sat_hist = cv.calcHist([img_hsv],[1],mask,[256],[0,256])
val_hist = cv.calcHist([img_hsv],[2],mask,[256],[0,256])
plt.plot(hue_hist)
plt.plot(sat_hist)
plt.plot(val_hist)
plt.legend(['hue','sat','value'])
plt.show()
h = np.arange(180, dtype=np.uint8).reshape(1, 180, 1)
s = (np.ones(180, dtype=np.uint8)*255).reshape(1, 180, 1)
v = s.copy()
hs = np.append(h, s, axis=2)
hsv = np.append(hs, v, axis=2)
hsv = np.repeat(hsv, 20, axis=0)
rgb = cv.cvtColor(hsv, cv.COLOR_HSV2RGB)

plt.figure(figsize=(6,3.2))
plt.imshow(rgb)
plt.yticks([])
# plt.xticks([0,20,30,40, 60, 120, 180], fontsize=15)
plt.tight_layout(pad=0.3)
plt.show()
np.sum(hue_hist[90:140])/np.sum(hue_hist)
hues_in_mask = img_hsv[:,:,0][mask == 255]
sats_in_mask = img_hsv[:,:,1][mask == 255]
```

```
vals_in_mask = img_hsv[:,:,2][mask == 255]
plt.hist2d(hues_in_mask.ravel(), sats_in_mask.ravel(), bins=(40, 20))
plt.xlim(0,179)
plt.ylim(0,255)
plt.xlabel('Hue')
plt.ylabel('Saturation')
plt.show()
plt.hist2d(hues_in_mask.ravel(), vals_in_mask.ravel(), bins=(40, 20))
plt.xlim(0,179)
plt.ylim(0,200)
plt.xlabel('Hue')
plt.ylabel('Value')
plt.show()
plt.hist2d(sats_in_mask.ravel(), vals_in_mask.ravel(), bins=(40, 20))
plt.xlim(0,255)
plt.ylim(0,200)
plt.xlabel('Saturation')
plt.ylabel('Value')
plt.show()
print('Average')
print('hue',np.mean(hues_in_mask))
print('sat',np.mean(sats_in_mask))
print('val',np.mean(vals_in_mask))

# %%

tiffs = glob(os.path.join('/directory/Alkal_Data/','*.tiff'))
tiffs

def analyze_tiff(tiff):
    img_fh = tiff
    results_file_prefix = os.path.basename(img_fh).removesuffix('.tiff')
    img = cv.imread(img_fh,cv.IMREAD_COLOR)
    img_hsv = cv.cvtColor(img, cv.COLOR_BGR2HSV)
    fig, ax = plt.subplots(3,1)
    # Plot original image
    ax[0].imshow(cv.cvtColor(img_hsv, cv.COLOR_HSV2RGB))
    mask = cv.inRange(img_hsv, np.array([0,0,0]), np.array([255,255,200]))
    # Plot mask
    ax[1].imshow(mask)
    res = cv.bitwise_and(img_hsv, img_hsv, mask=mask)
    # Plot maksed image
    ax[2].imshow(cv.cvtColor(res, cv.COLOR_HSV2RGB))
    fig.suptitle(results_file_prefix)
    plt.savefig('/directory/'+results_file_prefix+'masked_image.jpg')
    plt.close()

    # Plot hue histograms
    hue_hist = cv.calcHist([img_hsv],[0],mask,[256],[0,256])
    sat_hist = cv.calcHist([img_hsv],[1],mask,[256],[0,256])
    val_hist = cv.calcHist([img_hsv],[2],mask,[256],[0,256])
    fig, ((ax1, ax2), (ax3, ax4)) = plt.subplots(2, 2)
    ax1.plot(hue_hist)
    ax1.plot(sat_hist)
    ax1.plot(val_hist)
    ax1.legend(['hue','sat','value'])
```

```
    # Plot joint distributions
    hues_in_mask = img_hsv[:,:,0][mask == 255]
    sats_in_mask = img_hsv[:,:,1][mask == 255]
    vals_in_mask = img_hsv[:,:,2][mask == 255]

    hue_sat_hist2d, _,_,_ = ax2.hist2d(hues_in_mask.ravel(),
sats_in_mask.ravel(), bins=(40, 20), range=[[0,180],[0,255]])
    ax2.set_xlim(0,179)
    ax2.set_ylim(0,255)
    ax2.set_xlabel('Hue')
    ax2.set_ylabel('Saturation')

    hue_val_hist2d, _,_,_ = ax3.hist2d(hues_in_mask.ravel(),
vals_in_mask.ravel(), bins=(40, 20), range=[[0,180],[0,200]])
    ax3.set_xlim(0,179)
    ax3.set_ylim(0,200)
    ax3.set_xlabel('Hue')
    ax3.set_ylabel('Value')

    sat_val_hist2d, _,_,_ = ax4.hist2d(sats_in_mask.ravel(),
vals_in_mask.ravel(), bins=(40, 20), range=[[0,255],[0,200]])
    ax4.set_xlim(0,255)
    ax4.set_ylim(0,200)
    ax4.set_xlabel('Saturation')
    ax4.set_ylabel('Value')

    fig.suptitle(results_file_prefix)
    fig.tight_layout()
    plt.savefig('/directory/'+results_file_prefix+'300dpidistributions.jpg.',
dpi=300)
    plt.close()

    # Fraction blue pixels
    blue_pixels = np.sum(hue_hist[110:130])/np.sum(hue_hist)

    # Fraction redish pixels
    redish_pixels = np.sum(hue_hist[0:30])/np.sum(hue_hist)

    # Mean HSV
    mean_hue = np.mean(hues_in_mask)
    mean_sat = np.mean(sats_in_mask)
    mean_val = np.mean(vals_in_mask)

    return results_file_prefix, blue_pixels, redish_pixels, mean_hue,
mean_sat, mean_val, \
        hue_sat_hist2d, hue_val_hist2d, sat_val_hist2d

# Dictionary to hold results:
results_dict = {'Fish_ID':[], 'prop_blue_pixels':[],
'prop_redish_pixels':[],\
                'mean_hue':[], 'mean_sat':[], 'mean_val':[]}
pixel_dist_dict = {'hue_sat_hist2d':{'Wt':[], 'Het':[], 'Homo':[]}, \
                   'hue_val_hist2d':{'Wt':[], 'Het':[], 'Homo':[]}, \
                       'sat_val_hist2d':{'Wt':[], 'Het':[], 'Homo':[]}}

for tiff in tiffs:
```

```
    results_file_prefix, blue_pixels, redish_pixels, mean_hue, mean_sat,
mean_val, \
        hue_sat_hist2d, hue_val_hist2d, sat_val_hist2d = analyze_tiff(tiff)
    results_dict['Fish_ID'].append(results_file_prefix)
    results_dict['prop_blue_pixels'].append(blue_pixels)
    results_dict['prop_redish_pixels'].append(redish_pixels)
    results_dict['mean_hue'].append(mean_hue)
    results_dict['mean_sat'].append(mean_sat)
    results_dict['mean_val'].append(mean_val)
    results_df = pd.DataFrame.from_dict(results_dict)
    results_df
    # results_df.to_csv('/directory/results.csv')

    geno = results_file_prefix.split('_')[1]
    pixel_dist_dict['hue_sat_hist2d'][geno].append(hue_sat_hist2d)
    pixel_dist_dict['hue_val_hist2d'][geno].append(hue_val_hist2d)
    pixel_dist_dict['sat_val_hist2d'][geno].append(sat_val_hist2d)


genotype = [i.split('_')[1] for i in results_df['Fish_ID']]
results_df['genotype'] = genotype
results_df.head()

# Import necessary modules
from scipy import stats

# Plot results
order = ['Wt','Het', 'Homo']

# Perform one-way ANOVA test and post-hoc Tukey test on prop_blue_pixels data
F, p = stats.f_oneway(results_df[results_df['genotype'] ==
'Wt']['prop_blue_pixels'], results_df[results_df['genotype'] ==
'Het']['prop_blue_pixels'], results_df[results_df['genotype'] ==
'Homo']['prop_blue_pixels'])
if p < 0.05:
    from statsmodels.stats.multicomp import pairwise_tukeyhsd
    tukey = pairwise_tukeyhsd(endog=results_df['prop_blue_pixels'],
groups=results_df['genotype'], alpha=0.05)
    print(tukey.summary())

# Perform Mann Whitney U test on prop_blue_pixels and Prop_redish_pixels data
from scipy.stats import mannwhitneyu
homo_prop_blue = results_df[results_df['genotype']=='Wt']['prop_blue_pixels']
wt_prop_blue = results_df[results_df['genotype']=='Het']['prop_blue_pixels']
U1, p = mannwhitneyu(homo_prop_blue, wt_prop_blue, alternative='less')
print(p)

wt_prop_red = results_df[results_df['genotype']=='Wt']['prop_redish_pixels']
het_prop_red =
results_df[results_df['genotype']=='Het']['prop_redish_pixels']
U1, p = mannwhitneyu(wt_prop_red, het_prop_red, alternative='less')
print(p)

homo_prop_red =
results_df[results_df['genotype']=='Homo']['prop_redish_pixels']
het_prop_red =
results_df[results_df['genotype']=='Het']['prop_redish_pixels']
```

```
U1, p = mannwhitneyu(homo_prop_red, het_prop_red, alternative='greater')
print(p)

homo_prop_red =
results_df[results_df['genotype']=='Homo']['prop_redish_pixels']
wt_prop_red = results_df[results_df['genotype']=='Wt']['prop_redish_pixels']
U1, p = mannwhitneyu(homo_prop_red, wt_prop_red, alternative='greater')
print(p)


# Plot boxplots
sns.boxplot(x='genotype', y='prop_blue_pixels', data=results_df, order =
order, color="skyblue", fliersize=0)
sns.stripplot(x='genotype', y='prop_blue_pixels', data=results_df, order =
order, jitter=True, size=6, color='black')
plt.xlabel('Genotype')
plt.ylabel('Proportion Blue Pixels')
plt.title("Proportion Blue Pixels by Genotype")
plt.savefig('/directory/'+results_file_prefix+'BluishPixelBoxPlots.jpg.',
dpi=300)
plt.show()

# Plot boxplots
order = ['Wt','Het', 'Homo']
sns.boxplot(x='genotype', y='prop_redish_pixels', data=results_df,
order=order, color="crimson", fliersize=0)
sns.stripplot(x='genotype', y='prop_redish_pixels', data=results_df, order =
order, jitter=True, size=6, color='black')
plt.xlabel('Genotype')
plt.ylabel('Proportion Red Pixels')
plt.title("Proportion Red Pixels by Genotype")
plt.savefig('/directory/'+results_file_prefix+'RedishPixelBoxPlots.jpg.',
dpi=300)
plt.show()

distributions = ['hue_sat_hist2d']
genotypes = ['Wt', 'Het', 'Homo']

for d in distributions:
    # Set figure size to 10 inches wide and 12 inches tall
    fig, ax = plt.subplots(3,1, figsize=(12,10))
    for j,g in enumerate(genotypes):
        aggregate = np.zeros(pixel_dist_dict[d][g][0].shape)
        for i in pixel_dist_dict[d][g]:
            aggregate += i
        # Set x and y tick positions
        ax[j].set_xticks(np.arange(0, 181, 30))
        ax[j].set_yticks(np.arange(0, 256, 50))
        # Set x and y tick labels
        ax[j].set_xticklabels(['0', '30', '60', '90', '120', '150', '180'])
        ax[j].set_yticklabels(['0', '50', '100', '150', '200', '250'])
        ax[j].set_title("Hue vs Saturation", fontsize=20)
        ax[j].set_xlabel("Hue", fontsize=16, color="black")
        ax[j].set_ylabel("Saturation", fontsize=16, color="black")
        # Set the x and y axis limits
        ax[j].imshow(np.rot90(aggregate), extent=[0,180,0,255])
    fig.tight_layout()
```

```
    plt.savefig('/directory/'+d+'_joint-distributions.png', dpi=300)
    plt.show()

distributions = ['hue_val_hist2d']
genotypes = ['Wt', 'Het', 'Homo']

for d in distributions:
    # Set figure size to 10 inches wide and 12 inches tall
    fig, ax = plt.subplots(3,1, figsize=(12,10))
    for j,g in enumerate(genotypes):
        aggregate = np.zeros(pixel_dist_dict[d][g][0].shape)
        for i in pixel_dist_dict[d][g]:
            aggregate += i
        # Set x and y tick positions
        ax[j].set_xticks(np.arange(0, 181, 30))
        ax[j].set_yticks(np.arange(0, 201, 50))
        # Set x and y tick labels
        ax[j].set_xticklabels(['0', '30', '60', '90', '120', '150', '180'])
        ax[j].set_yticklabels(['0', '50', '100', '150', '200'])
        ax[j].set_title("Hue vs Value", fontsize=20)
        ax[j].set_xlabel("Hue", fontsize=16, color="black")
        ax[j].set_ylabel("Value", fontsize=16, color="black")
        # Set the x and y axis limits
        ax[j].imshow(np.rot90(aggregate), extent=[0,180,0,200])
    fig.tight_layout()
    plt.savefig('/directory/'+d+'_joint-distributions.png', dpi=300)
    plt.show()

distributions = ['sat_val_hist2d']
genotypes = ['Wt', 'Het', 'Homo']

for d in distributions:
    # Set figure size to 10 inches wide and 12 inches tall
    fig, ax = plt.subplots(3,1, figsize=(12,10))
    for j,g in enumerate(genotypes):
        aggregate = np.zeros(pixel_dist_dict[d][g][0].shape)
        for i in pixel_dist_dict[d][g]:
            aggregate += i
        # Set x and y tick positions
        ax[j].set_xticks(np.arange(0, 256, 50))
        ax[j].set_yticks(np.arange(0, 201, 50))
        # Set x and y tick labels
        ax[j].set_xticklabels(['0', '50', '100', '150', '200', '250'])
        ax[j].set_yticklabels(['0', '50', '100', '150', '200'])
        ax[j].set_title("Saturation vs Value", fontsize=20)
        ax[j].set_xlabel("Saturation", fontsize=16, color="black")
        ax[j].set_ylabel("Value", fontsize=16, color="black")
        # Set the x and y axis limits
        ax[j].imshow(np.rot90(aggregate), extent=[0,255,0,200])
    fig.tight_layout()
    plt.savefig('/directory/'+d+'_joint-distributions.png', dpi=300)
    plt.show()
```
